# Morphodynamics of chloroplast network control light-avoidance response in the non-motile dinoflagellate *Pyrocystis lunula*

**DOI:** 10.1101/2024.04.30.591832

**Authors:** Nico Schramma, Gloria Casas Canales, Maziyar Jalaal

## Abstract

Photosynthetic algae play a significant role in oceanic carbon capture. Their performance, however, is constantly challenged by fluctuations in environmental light conditions. Here, we show that the non-motile single-celled marine dinoflagellate *Pyrocystis lunula* can internally contract its chloroplast network in response to light. By exposing the cell to various physiological light conditions and applying temporal illumination sequences, we find that network morphodynamics follows simple rules, as established in a mathematical model. Our analysis of the chloroplast structure reveals that its unusual reticulated morphology constitutes properties similar to auxetic metamaterials, facilitating drastic deformations for light-avoidance, while confined by the cell wall. Our study shows how the topologically complex network of chloroplasts is crucial in supporting the dinoflagellate’s adaptation to varying light conditions, thereby facilitating essential life-sustaining processes.

---

Light is fundamental for life, yet excessive exposure can be detrimental for biological processes. Photosynthetic organisms have developed multiple strategies across scales to cope with fluctuations of light, from molecular adaptation responses such as Non-Photochemical Quenching (NPQ) to the biased growth towards light (phototropism) on organismal scale [1–3]. Photoadaptation in the form of increased thermal dissipation of photo-excited chlorophyll, which displays a major component of NPQ, occurs within tens of seconds to minutes [4], while plant tropism takes longer time scales of hours to days [3, 5]. Another photoadaptation mechanism, at a single-cell level, includes the motion and rearrangement of chloroplasts to optimize light absorption or to avoid photo-damage [2, 6–10]. Such strategies are fundamental for survival in fluctuating environments, displaying interesting features such as computation and integration in the context of plant phototropism [11–13], and dynamical phase transitions of chloroplast motion [10]. In land plants, the collective motion of individually sensing and moving disk-shaped chloroplasts [9, 14] leads to large-scale re-arrangements inside leaf cells, aiming to optimize for light uptake while avoiding strong light. However, a vast amount of photosynthesis happens in aquatic environments, especially the oceans [15–17]. Some single-celled algae can actively swim away from or towards light [18–21] and even less complex organisms such as cyanobacteria, like *Trichodesmium* [22] and *Synechocystis* [23, 24], can move collectively or individually as a response to light. Non-motile photosynthetic organisms, nonetheless, have adopted different strategies by moving their chloroplasts within their cell bodies [25, 26].

Here, we study the light adaptation strategy of a non-motile marine dinoflagellate. Dinoflagellates are a large and distinct group of photosynthetic and mixotrophic algae [27, 28] that, unlike most other photosynthetic organisms, underwent tertiary endosymbiosis by engulfing algae containing a secondary plastid [29–32]. This process led to chloroplasts with three membranes [33]. Mostly known for their bioluminescence [34–36], the non-motile, ∼ 50 − 100 *µ*m-sized dinoflagellate *Pyrocystis lunula*, native to warm waters worldwide, can reorganize its internal architecture to switch from a photosynthetic day-phase to a bioluminescent night-phase. This reorganization is orchestrated by actively moving their chloroplasts and bioluminescent organelles (scintillons) towards and away from the cell center, following a circadian rhythm [37–39]. Interestingly, it has been observed that the same intracellular motion can be triggered by strong light [40], leading to a compression of the chloroplast within just a couple of minutes, without deformation of the thick cell wall [41]. This observation raises the question of how this drastic intracellular rearrangement can be coordinated under confinement and within such a short time frame. Here, we study the dynamics of chloroplast motion as a light adaptation mechanism in *P. lunula* and report that it features an active reticulated chloroplast network, capable of fast morphological changes. This enables the organelle to undergo significant deformations for efficient photo-avoidance motion in response to environmental changes.

## Cytoplasmic space contracts under strong light

We studied the adaptation of the chloroplast area of individual *Pyrocystis lunula* cells to white-light exposure under physiologically relevant light conditions of its natural habitat at depth of 60 − 100 m [42] (Supplementary Text Section I). Upon white-light stimulation, chloroplasts move towards the cytoplasmic core area within approximately 10 min (Fig. 1A) [40]. During this process, the relative absorbed light (Fig. 1C) and the projected area *𝒜* of the chloroplast (Fig. 1D) exhibit similar dynamics for various light irradiances ranging from 2.8 to 41.4 mW*/*cm^2^ (fig. S1, Movie S1). After a short time of approximately 1 − 2 min, characterized by a positive curvature, both the projected area and the absorbed light follow an exponential decay, eventually reaching saturation at levels dependent on the light intensity. These measurements are correlated: as the chloroplast retracts, more light can pass through the cell, leading to reduced absorption of up to 10 %.

**Fig. 1:**
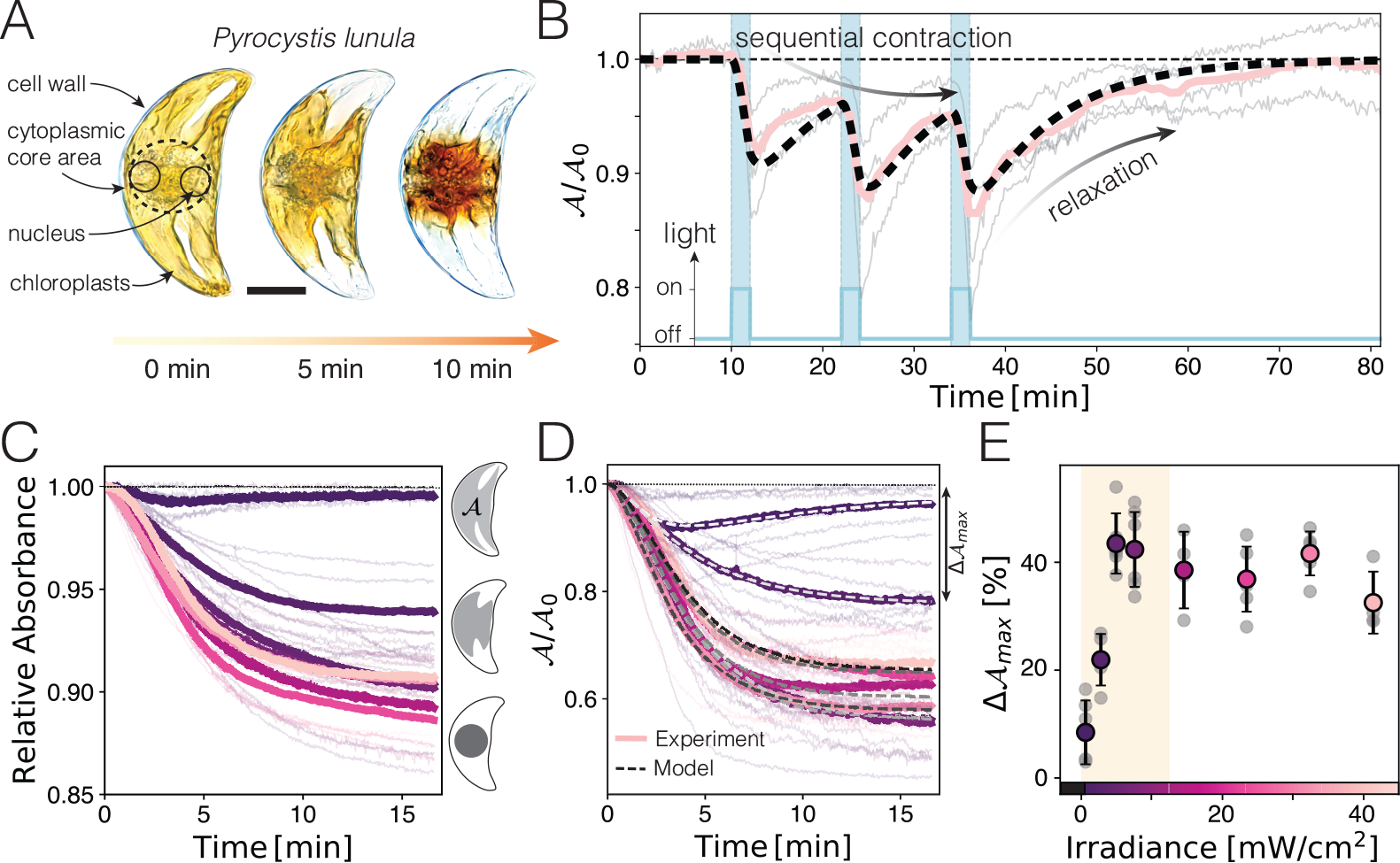
Chloroplast area of *Pyrocystis lunula* undergoes rapid changes under strong light illumination. (**A**) Upon light activation, the chloroplast (yellow) contracts and compacts (darker color), achieving significant change in less than 10 min. Scalebar: 20 *µ*m. (**B**) Periodic light stimulation (blue curve), causes the chloroplast area *𝒜* (normalized by initial area *𝒜*_0_, *N* = 4, orange curve: mean) to decrease under strong light and expand under dim light. The dynamics are well described by an active viscoelastic model (dashed line). (**C** and **D**) Relative absorbance and area decline in response to varying light intensities in specimens initially adapted to dim light, indicating increased light transmission in the light-avoidant state. Higher light intensities cause a stronger response (colorscale in **E**). (**E**) The maximum area reduction correlates with light irradiance. Beyond physiologically tractable light conditions (yellow box) the contraction response saturates to a maximal decrease of the chloroplast area by 40 %.

Contrary to the contraction scenario described above, exposure to dim white light (0.6 mW*/*cm^2^) triggers a transient response wherein the chloroplast initially contracts towards the cell center but eventually expands again (Fig. 1D). These observations share similarities with terrestrial plants’ transient photo-response at the intermediate light intensity [43, 44]. Large stimulation-irradiances, which exceed both the estimated ecologically relevant light conditions of 0.2 − 12 mW*/*cm^2^ (Supplementary Text Section I) and the culturing conditions *I* = 0.27 mW*/*cm^2^ (Materials and Methods) lead to maximal contraction of Δ*𝒜*_*ax*_ ≈ 40 %, corresponding to the size of the cytoplasmic core area (Fig. 1A,E).

### Dynamical testing and mathematical model of chloroplast contraction

Alternating white- (on) and red-light (off) irradiation controls the chloroplast contraction and relaxation dynamics towards and away from the cytoplasmic core area (Fig. 1B, Movie S2,S3): the chloroplast network rapidly responds to white-light and slowly relaxes upon red-light exposure. Long-time illumination with weak red light (0.2 mW*/*cm^2^) leads to a complete expansion of the chloroplasts over the entire cell (Fig. 1B, Time*>* 50 min). The dynamical tests verify a mathematical model (dotted lines Fig. 1B,D, 3G, 4C,D) (Materials and Methods, Supplementary Text Section II), based on two chemical species that control a light-avoidant and a light-accumulation response (contraction or expansion), respectively (fig. S2).

The model enables the extraction of two relevant chemical signaling timescales, *τ*_1_ and *τ*_2_, along with a timescale for active contraction *τ*_*KV*_ and expansion 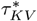. We find that the chemical signaling of a photo-avoidance response occurs at a time scale of *τ*_1_ = 1.8 ±0.5 min (mean ± SD, *N* = 44), while a photo-attractive signal leading to a transient response is noted at *τ*_2_ = 2.5 ± 0.9 min (*N* = 6). The timescales *τ*_*KV*_ and 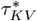 emerge from the dynamics of the driving mechanism, relying on actin, microtubules, and molecular motors such as myosin [39] (also see Supplementary Text Section III for inhibitory treatment), and found to be *τ*_*KV*_ = 2.7 ± 0.9 min (*N* = 44) for contraction and 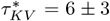 min (*N* = 6) for expansion of the chloroplast. Notably, our model can be interpreted as a filter for environmental stimuli (Supplementary Text Section II), effectively filtering out high-frequency noise (with short time scales) such as the light fluctuations of surface waves (Supplementary Text Section I).

### Topologically complex chloroplast network allows efficient contraction

The contraction of the projected area reaches up to 40 % of the cell area, necessitating a remarkably large deformation of the cytoplasmic material. Such deformations within hard confinement, in this case, constituted by the cell wall of *P. lunula* [41], are subject to physical constraints. To elucidate these constraints, we draw analogies to the compression of fluids and solids: an incompressible fluid cannot “contract” uniformly towards the center under confinement, as its volume must be conserved. Similarly, an elastic solid, when compressed from one direction, will expand in orthogonal directions, a behavior characterized by a positive Poisson ratio (*ν* = 0.5 for isotropic incompressible elastic materials in three dimensions). Under confinement, this property (*ν >* 0) leads to effective strain-stiffening, as the expansion into orthogonal directions is prevented and therefore internal stresses redirect. However, structured metamaterials defy positive Poisson ratios by allowing non-linear deformations such as buckling or the action of rotating hinges [45, 46]. Auxetic behavior, characterized by *ν <* 0, naturally arises in polymer networks [47], foams [48], and poro-elastic materials such as cork (*ν* ≈ 0) [49], enabling uni- or multi-directional contraction, and thus facilitating efficient deformations even within confined spaces.

Using confocal auto-fluorescence imaging of the chloroplasts (Materials and Methods) we uncover the chloroplast reticulum (Fig. 2A). This intricate network structure shows similarities to observations made in the context of spontaneous diurnal chloroplast relocations [38, 39] and rapid changes of buoyancy in the related species *Pyrocystis noctiluca* [50]. Continuous blue light stimulation (*λ* = 470 ± 50 nm) of the cell, induces contraction of the chloroplast network over time. We uncover two mechanisms which choreograph this chloroplast photo-adaptaion motion: cytoplasmic strands move towards the center while they simultaneously contract in a manner reminiscent of buckling (Fig. 2A, Movie S4-S6). Buckling, or inward-folding, allows the structure to compact into the space between the cytoplasmic strands (*ν* ≲ 0), circumventing strain-stiffening typically expected from uniform bulk contractions under confinement (*ν >* 0). Moreover, we observe a notable thickening of some strands during contraction, indicating a flow of material within the structure (Fig. 2A, Movie S4-S6). Under dim red light, the chloroplast expands again over a longer time scale (Fig. 2B, Movie S6). Although the chloroplast network appears slightly different before and after the contraction and expansion, the main geometrical features remain unchanged: when fully spread, the network is characterized by a few very long cytoplasmic strands (Fig. 2C), which extend outwards along the cell’s long body axis (Fig. 2D). During contraction, these distinctive features are lost as the chloroplast reticulum obtains a more spherical shape. Remarkably, despite these morphological changes, the network topology remains largely unchanged before contraction and after expansion, suggesting a permanent connection among nodes without dynamic rewiring of the network (Fig. 2E). The apparent loss of holes, as indicated by the decline in Genus, can be attributed to the increased contact between strands, which complicates the identification of individual strands.

**Fig. 2:**
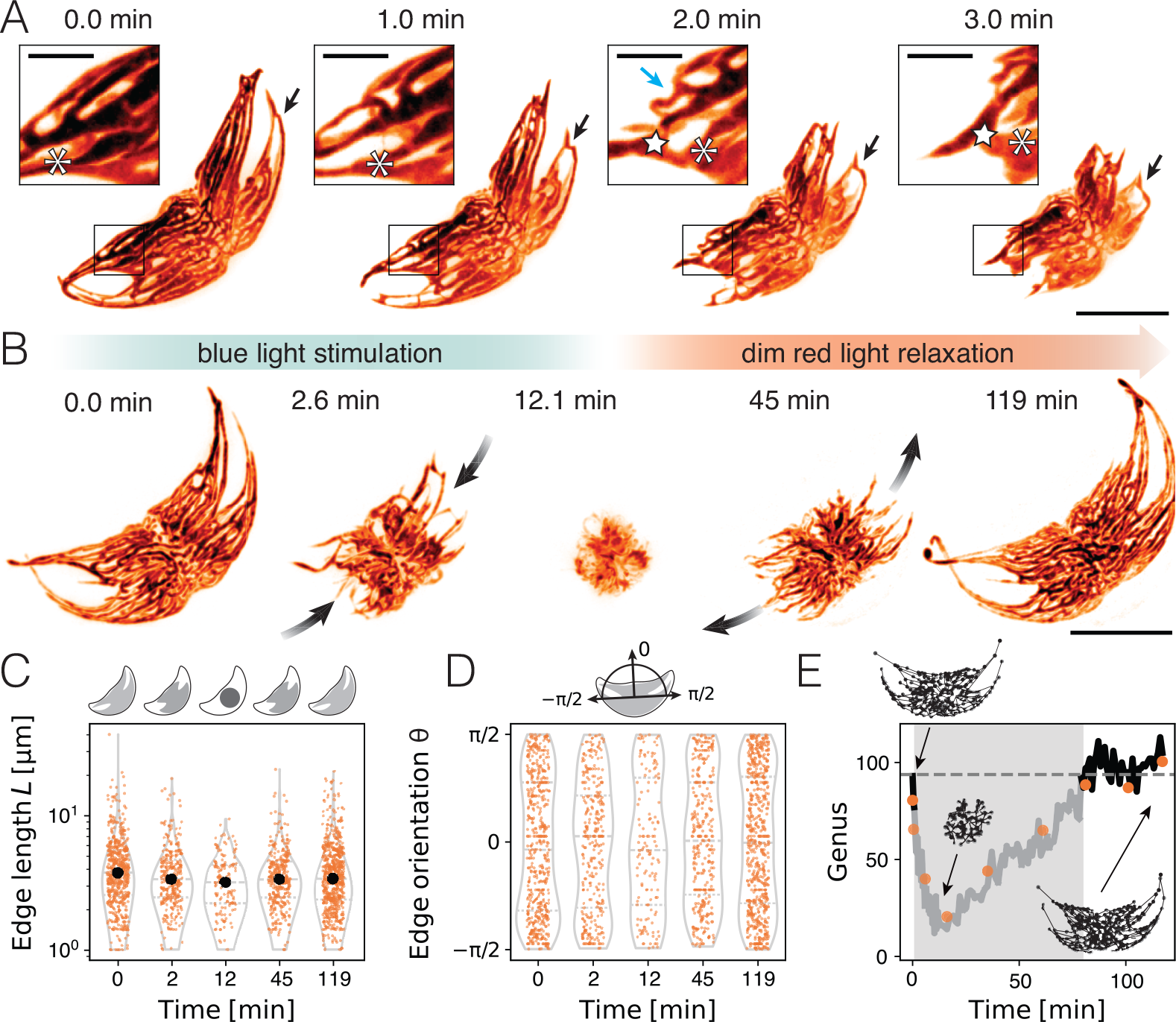
Dynamic photo-avoidance and photo-accumulation of the chloroplast reticulum. Autofluorescence imaging of chloroplasts reveals reticulated structure. (**A**) Light stimulation beginning at *t* = 0 min causes the chloroplast network to contract, by moving strands towards the cells’ center and deforming them (arrows). A compact structure is achieved by reducing hole sizes of the network. Scale bar: 40 *µ*m. Inset: Examining an area of dynamical deformation of a strand of the chloroplast network, two nodes (asterisk and star) change their position as well as their distance from one another. The blue arrow indicates a deforming cytoplasmic strand leading to the closure of a hole in the network. Scale bar: 10 *µ*m. (**B**) Chloroplast contraction under blue light stimulation and subsequent expansion under the dim red light environment. (**C**) Edge length statistics of the chloroplast network (from B) over time. Large edges shorten and extend again. (**D**) Distribution of orientation of edges. Orientations align with the major axis of the cell, but during contraction, the distribution flattens. (**E**) Topological measurements over time, indicating similarity of structures before and after the stimulation. Black line: Genus estimated by Betti number *β*_1_ of the network. Orange points: Genus measured from Euler characteristic *χ* of the mask image. During contraction chloroplast strands touch and seemingly close holes leading to an apparent decrease in detected genus (gray area). Genus before contraction and after expansion has similar values.

We analyze the spatiotemporal dynamics of the networks’ nodes over time (Fig. 3A,B,E, Materials and Methods, Movie S7) to quantify the contraction process. Our findings reveal a wide distribution in node speeds, which vary over time (Fig. 3A,C). Predominantly, nodes close to the cytoplasmic core area exhibit slow, diffusive motion characterized by a time-linear mean-squared displacement (MSD ∝ *τ*). Conversely, nodes located farther from the cell center move ballistically (MSD ∝ *τ* ^2^) (Fig. 3D). Thus, longer strands, initially located far from the cytoplasmic core area, move rapidly towards the center (Fig. 3E,H), while reducing their length, resulting in the reduction of chloroplast’s 2D-projected area 𝒜 (Fig. 3G). Interestingly, the strands follow along their initial configuration, as indicated by the overlay of trajectories on the network’s 2D projection at *t* = 0 min in figure 3A,E. This may reflect a signature of their actin- and microtubule-mediated driving mechanism [39] and a co-alignment to those networks. Further analysis of the network, demonstrates that the speed of the nodes scales with their distance from the center, *v* ∝ *D*, aligning with the expected behavior from the one-dimensional contraction of an elastic material under a constant strain rate (Fig. 3H, inset). This observation confirms that the complex topology of the network enables the material to contract, dominantly, along one dimension in a manner similar to an elastic material (*ν* ∼ 0). These findings have been valuable in developing the mathematical model (Supplementary Text Section II).

**Fig. 3:**
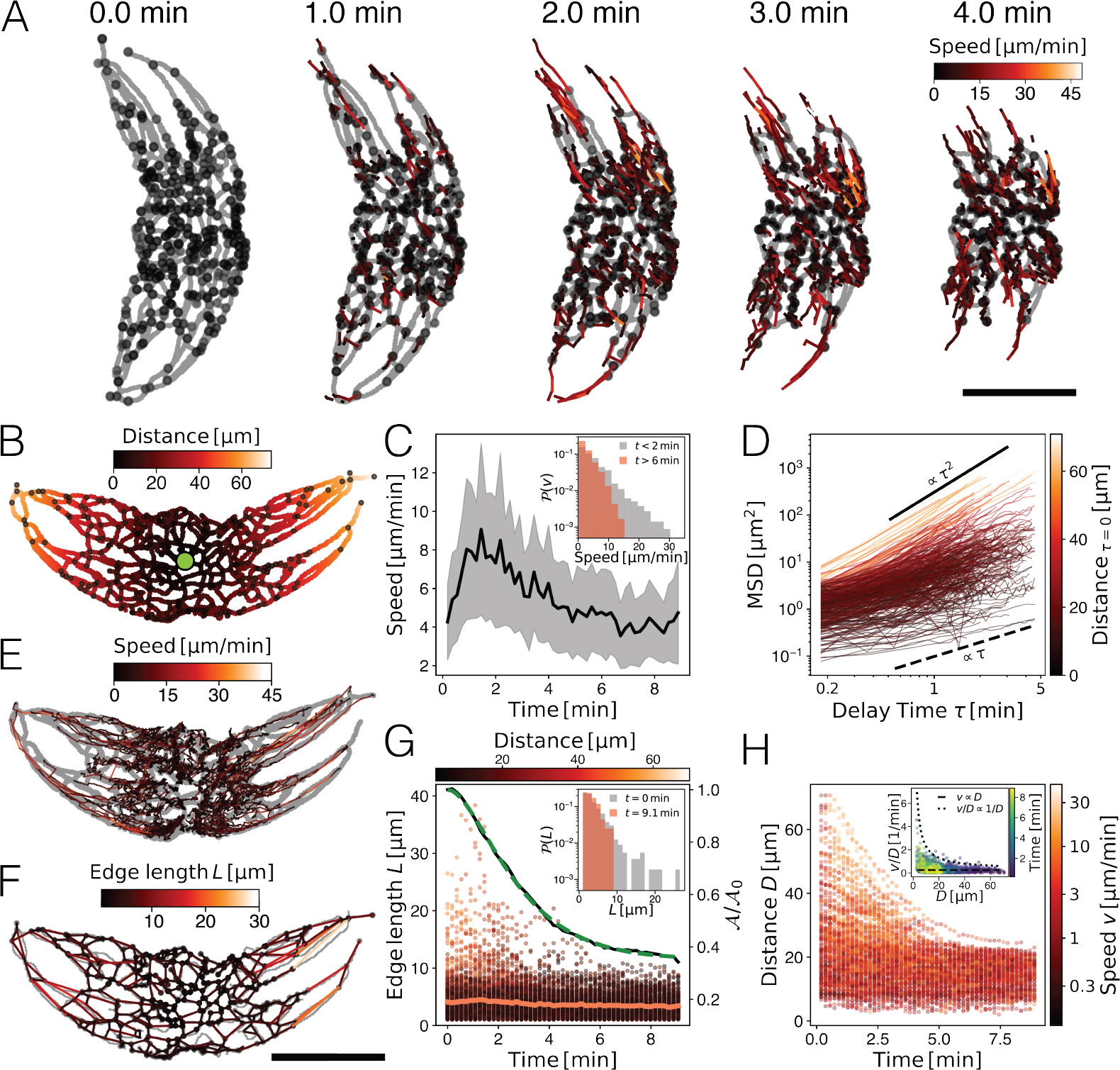
Network dynamics of the chloroplast reticulum. (**A**) Time series of chloroplast contraction dynamics. The nodes (black dots) of the underlying three-dimensional network move on paths inward (color: speed). Gray: 2d projection of the underlying skeleton. (**B**) Distance-map from the center point (green dot) on a 2d-projected mask of the experiment, as in (A) at *t* = 0 min.(**C**) Average individual node speed (black line: mean *±* SD) increases during contraction and subsequently decreases. Inset: the speed distribution is heavy-tailed. Initially fast trajectories (*t <* 2 min) slow down at long times *t >* 6 min. (**D**) Mean-squared displacement (MSD) of individual trajectories ranges from diffusive trajectories (MSD *∝ τ*) in the center (colors corresponding to the distance from the center (B) at the beginning of the trajectory) to purely ballistic trajectories (MSD *∝ τ* ^2^) originating at the periphery. (**E**) Trajectories are mapped on the skeleton of the mask (gray) and colored according to their local speed reaching up to 30 *µ*m*/*min. (**F**) Network representation of the chloroplast. Edges are colored by length. The network closely represents the skeleton of the mask (gray). (**G**) The distribution of edge lengths of the network decays over time. The mean edge length (orange line) is hardly affected. Colorbar: edge distance to the center corresponding colors in (B) and (D). The shrinkage of the longest edges follows a similar trend to the decrease in the relative projected area of the chloroplast network (black line, right axis). The model fit to the projected area curve (dashed green line) shows good agreement with the experimental observations. Inset: The network has initially (*t* = 0 min) a few very long trajectories (grey), which disappear at long times (red). (**H**) Correlation of distance and speed. Far-distanced nodes travel at higher speeds (brighter colors) as compared to centrally located nodes. Note the logarithmic colorbar. Inset: “Strain rate” (speed over distance) of every node is constant at large distances. The solid line corresponds to *v ∝ D* which is expected for elastic deformation at a constant strain rate. The dotted line corresponds to constant *v*. Colormap represents time. All scale bars: 40 *µ*m

### Chloroplast contraction shows local and global responses to local stimulation

To further shine light on the sensing mechanism, we locally stimulate the cell (Materials and Methods) with a blue low-power laser (*λ* = 488 nm, 3.25 mW*/*cm^2^). Although this power level may seem weak compared to the full-spectrum irradiance of white-light stimulation used in other experiments, it closely matches the optimal absorption peak of phototropins [51]. Phototropins, crucial photosensors for chloroplast positioning in terrestrial plants [9, 52, 53], are also present in *Pyrocystis lunula* [54], potentially accounting for the observed strong response.

We illuminate a 6.8 × 6.8 *µ*m^2^ region at both the center and periphery of the algae, respectively (Fig. 4A,B, Movies S8-S10). Peripheral stimulation leads to a localized contraction of one side of the chloroplast network towards the cytoplasmic core area. The opposite side of the chloroplast network moves less pronounced, but within the temporal resolution limit of 20 − 40 s. This rapid onset suggests that a fast diffusing signal triggers a long-range response across the cell of *L* ≈ 80 − 100 *µ*m within *τ* ≈ 20 − 40 s after light-stimulation begins, implying a diffusion coefficient on the order of *D* ≈ *L*^2^*/τ* ≈ 160 − 500 *µ*m^2^*/*s. Interestingly, the chloroplast retraction on the not-stimulated region is transient or less pronounced, as observed in small light intensity stimulation (Fig. 1D, Movie S8,S9), suggesting that the transmitted signal is depleted, as expected for diffusive signaling.

**Fig. 4:**
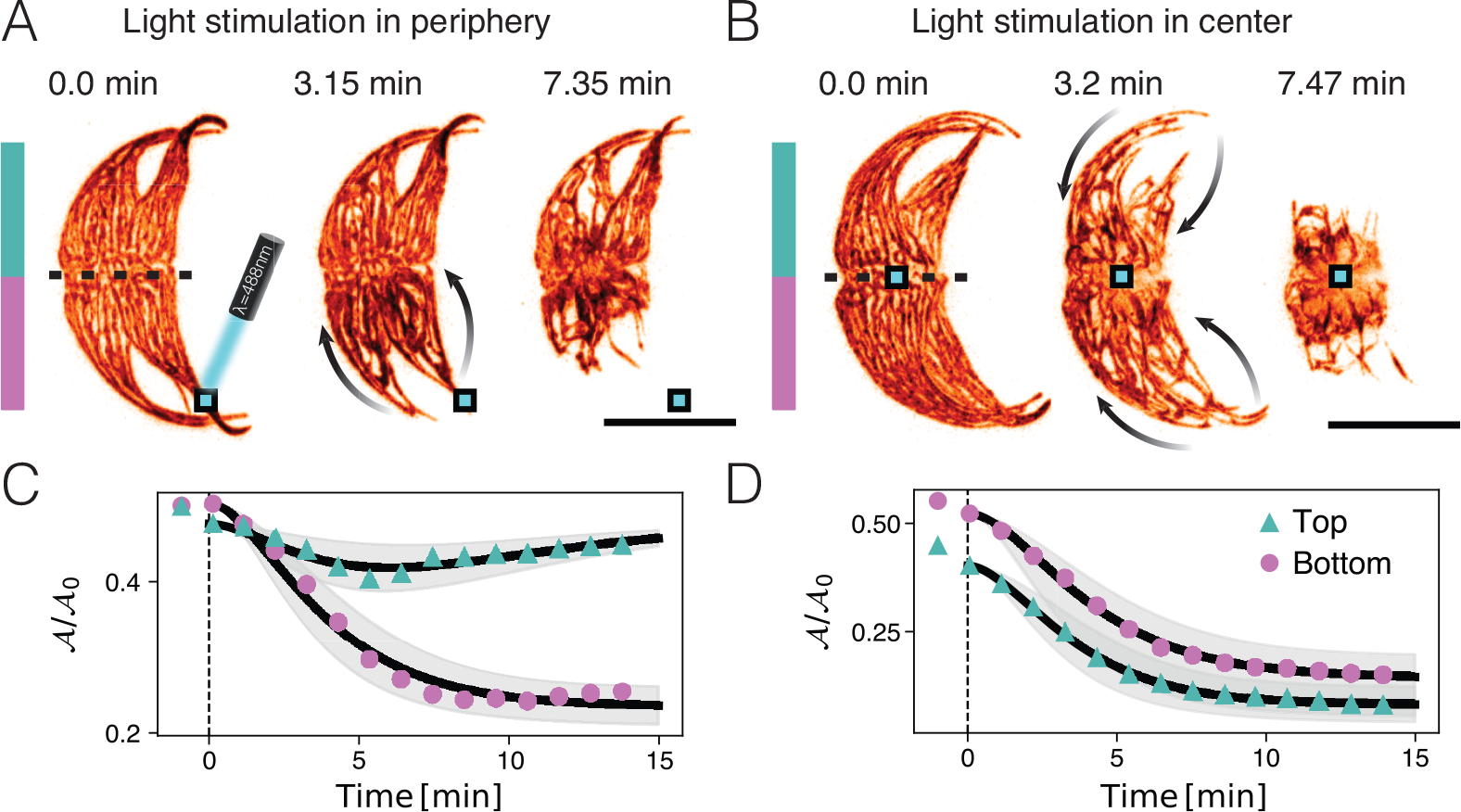
Local sensing leads to local or global response. (**A**) Local stimulation in the periphery with blue light (*λ* = 488 nm, *I* = 3.25 mW*/*cm^2^) leads to a one-sided response away from the stimulation region within a few minutes. Note the contraction of the network in the opposite site. Blue square indicates stimulation region. (**B**) Light stimulus in the center (blue square) leads to the global movement of the chloroplast network towards the center (arrows). (**C**) Area contraction in (A) upper and lower half of the cell (divided by dashed line in (A)). The lower part contracts rapidly, while the upper half undergoes a transient response. (**D**) Area contraction in (B). Upper and lower half of the cell undergo a symmetric response. See Movies S8-S10. Scale bar: 40 *µ*m.

Surprisingly, upon central stimulation in the cytoplasmic core area, all chloroplast strands contract towards the cell center, within similar time scales (Fig. 1B,D, Fig. 3A,G). This outcome reveals a symmetry-broken response: under peripheral stimulation, the chloroplast network contracts to move *away* from the light source, whereas central stimulation induces movement *towards* it. This pattern suggests that the chloroplast network’s contraction mechanism is inherently directed inward in response to strong light, regardless of the signal’s location.

## Discussion

In our experiments, we show that the chloroplasts of *Pyrocystis lunula* retract towards the cell’s center under strong white or blue light conditions and expand under weak red light conditions. This bidirectional movement of chloroplasts towards and away from light, mirrors the chloroplast photo-relocation motion seen in leaves of green plants [2, 3, 6, 9], nonetheless, using a fundamentally different dynamics. The significant light-induced retraction of chloroplasts leads to increased light transmission through the cells (Fig. 1C), suggesting that, similar to green plants, *Pyrocystis lunula* employs this mechanism as a means of light avoidance [2, 6, 7, 9]. Notably, under low white light conditions, we observe a transient response in the chloroplasts – initial fast contraction followed by slow expansion – suggesting a timescale-separated competition between these processes. This shows similarities with the observed counteraction of phototropin 1 and 2-mediated chloroplast motion [43, 55]. In fact, the transcriptome of *P. lunula* bears various phototropin 1 and phototropin 2-like sequences and LOV domains [54], pointing towards potential similarities in light sensation. The similarity of this organism’s light response to green plants is surprising, as the origin of chloroplasts in dinoflagellates is very distinct [27, 32].

At light intensities exceeding the natural physiological conditions of dinoflagellates, chloroplasts contract fully towards the cell center. Under such extreme conditions, the crowding of the chloroplast strands poses a mechanical limit for contraction, and consequently for this photo-avoidance mechanism. However, within their native physiological light conditions, *Pyrocystis lunula* responds via light-dependent chloroplast compaction, indicating a gradual adaptation response to various light conditions. Furthermore, we find that the relaxation phase following strong light-induced contraction occurs over a longer time scale, suggesting a different driving mechanism for the expansion of the chloroplasts.

The large-scale transport of organelles observed likely depends on the coordinated action of the actin and mi-crotubule networks, together with molecular motors such as myosin, as has been demonstrated in the context of the diurnal intracellular reorganization between the photosynthetic phase during the day and bioluminescent phase at night [39], in line with pharmacological perturbation experiments for light-adaptation (Supplementary Text III, fig. S3) in which we confirm that actin is necessary for bi-directional chloroplast motion. However, the exact driving mechanism of the chloroplast relocation in *P. lunula* remains unidentified and may differ from that of green plants, where chloroplast movement is primarily driven by the assembly of short actin (cp-actin) filaments and the transmission of polymerization forces towards the plasma membrane [9, 14, 55].

We elucidate the details of the reticulated morphology of chloroplasts and show that such a structural “design” offers mechanical advantages. Structured metamaterials, like this chloroplast morphology, facilitate buckling and other complex deformations [45, 46], enabling efficient chloroplast contraction under the confinement of the cell wall. *Pyrocystis noctiluca*, another species within the *Pyrocystis* genus, was found to have a reticulated cytoplasm, assisting with the vertical migration [50], showing another intricate link between the morphology of cytoplasmic space and its function in dinoflagellates.

We also showed that the dynamics of chloroplast contraction can be effectively modeled using a “visco-elastic” framework with chemically controlled stress applications. Although our coarse-grained model does not pinpoint the precise origins of observed elasticity and viscosity – whether from passive or active cellular components – it importantly enabled us to identify adaptation time scales and compare them with the ecologically relevant fluctuations. These time scales suggest that chloroplast motion serves as a feasible light adaptation strategy for environmental light variations persisting longer than 3 − 5 min. Indeed, such fluctuations notably exceed the duration of second-long light changes induced by waves but are in line with the motion of clouds obscuring the sun [56] and might complement NPQ.

Our experiments have also shown that chloroplast contraction is driven by local sensing, with directed relocation observed when chloroplasts are locally stimulated (Fig. 4). However, even though locally stimulated, the chloroplast network on the opposite side of the cell contract within 20 − 40 s, indicating a long-ranged signal transfer via fast diffusive signals (*D* ≈ 160 − 500 *µ*m^2^*/*s), such as calcium, recognized for its important roles in photosensory downstream signaling [53] and in *P. lunula* bioluminescence [57]. Interestingly, stimulation in the cytoplasmic core prompts chloroplasts to move towards rather than away from the center. This counter-intuitive behavior might result from the network’s topologically conserved structure and a photo movement that is inherently biased towards the cell center. This hypothesis, however, needs further investigation. Moreover, the intricate relationship between the chloroplast and nucleus, with the latter hosting a significant portion of the chloroplast genome [2, 58], suggests that chloroplast movement towards the nucleus in the cytoplasmic core [38] could also serve a photoprotective function, shielding genetic material from intense light damage. Our study provides the first evidence for such a mechanism in dinoflagellates, however, a comprehensive examination of the chloroplast-nucleus relationship is necessary to fully understand these dynamics.

Overall, the complex relationship between the geometry and topology of chloroplast structure and its dynamics provides a fertile ground for exploring intriguing physical dynamics with significant physiological implications, in the context of light-life interactions.

## Acknowledgements

We thank Ronald Breedijk for help with confocal imaging and localized light stimulation. We thank Joachim Goedhart, Mark Hink, Yuri Z. Sinzato, Jonas Veenstra, Corentin Coulais and Friedrich Kleiner for fruitful discussions. Confocal imaging was supported by the van Leeuwenhoek Centre for Advanced Microscopy, Section Molecular Cytology, Swammerdam Institute for Life Sciences, University of Amsterdam, a EuroBioImaging-node. MJ acknowledges the ERC grant no. “2023-StG-101117025, FluMAB”.

## Author Contribution

Nico Schramma: Conceptualization (lead); Investigation (equal); Validation (equal); Writing – original draft (lead); Visualization (lead); Formal analysis (lead); Writing – review and editing (equal); Software (lead); Methodology (lead)

Gloria Casas Canales: Conceptualization (supporting); Investigation (equal); Validation (equal) ;Writing – original draft (supporting); Visualization (supporting); Writing – review and editing (equal)

Maziyar Jalaal: Conceptualization (supporting); Writing – original draft (supporting); Visualization (supporting); Writing – review and editing (equal); Methodology (supporting); Supervision (lead); Funding (lead)

## Competing interests

The authors declare no competing interests.

## Data and materials availability

Data is available upon request at the corresponding author.

## APPENDIX

### I. MATERIALS AND METHODS

#### Cell culture

*Pyrocystis lunula* (Schütt) is cultured in *f/*2 medium in an incubator (Memmert) at 20 °C and a 12 : 12 day-night cycle with light irradiance at 0.27 mW*/*cm^2^.

#### Brightfield microscopy

We perform bright-field microscopy with a Nikon TI2 E microscope. Images are acquired using a Photometrics BSI Express camera at a frame rate of 1 − 2 frame*/*s. We use a 40*x/*1.2NA PLAN Apochromat objective and simultaneously measure 3 − 5 different positions using a motorized xy-stage totaling 4 − 8 cells for every light intensity setting (0.6, 2.8, 4.9, 7.6, 14.5, 23.4, 32.4, 41.4 mW*/*cm^2^). To allow for chloroplast expansion, imaging was performed using a red filter with a cut-on wavelength of *λ*_*c*_ = 625 nm, at light intensities of 0.1−0.4 mW*/*cm^2^. Periodic white (7.6 mW*/*cm^2^) and red light (0.1 mW*/*cm^2^) stimulation was controlled through a custom script in microManager [59].

#### Confocal microscopy

Global simulation experiments are performed with Nikon Ti Eclipse microscope equipped with a 60×/1.49NA or 40×/1.3NA oil objective, a confocal spinning disk unit (Yokogawa CSU-X1) with a microlense array, and an Andor iXonEM+ 897 electron-multiplying charged-coupled device (EMCCD) camera. Chlorophyll autofluorescence is stimulated at 640 nm wavelength while the emission bandpass filter (680 − 740 nm) is used. By using a piezo z-stage we record 50 − 120 z-steps (step size 0.3 − 1 *µ*m) within 11 − 20, s with a 30 − 60 ms excitation time. The brightfield path of the microscope was equipped with a 470 ± 50 nm blue filter to stimulate the cell at an intensity of 0.6 mW*/*cm^2^.

#### Local stimulation

Local stimulation experiments are performed using a 63× PLAN APO IR objective on a Nikon TI body with Nikon A1 resonant scanner. Imaging is performed using a 637nm laser line and emission filter at 650 nm. A whole z-stack consists of 80-100 steps of 0.5 *µ*m size. The pinhole diameter is fixed to 1.2 Airy disk diameters. Local stimulation is performed by applying a 488 nm laser at low irradiance 3.25 mW*/*cm^2^ in a small scanning region of 6.8×6.8 *µ*m^2^. One z-stack takes between 45 s and 60 s and the stimulation time is 3.9 s for center-stimuli or 7.8 s for peripheral stimuli.

#### Image processing

Image analysis was performed in Fiji [60] and Python using Napari [61] and scikit-image [62]. In the following paragraphs we will outline the different image processing steps for the different data we acquired.

#### Measuring chloroplast area from brightfield data

We customized a Fiji macro, which measures bright-field intensity in a hand-annotated region of interest (ROI) for sequential stimulation experiments, or generated ROI for constant illumination experiments. The ROI is generated by Gaussian smoothing of the first frame with a 21*px* kernel and subsequent thresholding using a triangle filter. Morphological closure with a 51*px*-diameter disk-shaped mask helps filling holes within the mask. To measure the area we calculate a the mean value of the background, subtract it by one standard deviation and employ it as a threshold value to discriminate the chloroplast material from the background light transmitted through the organism.

#### Analyzing 3D confocal data

We segment and skeletonize 4d stacks (x,y,z,t) by using a script developed within Napari. The following steps are performed for all time steps. First, we estimate the loss of light intensity deeper within the sample by fitting the Beer-Lambert law *I*(*z*) = *I*_0_*e*^*−z/λ*^ and correcting the z-stack accordingly. Next, we blur the x-y plane with a 1 px-wide Gaussian and subsequently use a top-hat filter with a (2.8, 2.8, 1) *µ*m kernel. Then, we use the triangle method to threshold and perform binary closing with a (1, 1, 1) *µ*m kernel. We label the obtained mask with a connected-component method and reject small labels *<* 20000 px. Three-dimensional graph analysis was performed after skeletonizing the label image [63] and generating a NetworkX graph [64]. Some closely located nodes of the graph are merging over time, hence we coarse-grain nodes that are less than 5*px* away from each other by deleting them and setting one node in the middle between them. Similarly, we reject end-nodes that have a single edge less than 9*px* long. The nodes of the graph represent the junctions of the chloroplast reticulum and are tracked using a nearest-velocity tracker in trackpy [65] with a (6 *µ*m, 6 *µ*m, 2 *µ*m) search window and 4-time step memory (≈ 40 − 60 s), as well as an adaptive search window to down to 10 % of the initial search window size, to enhance the trajectory matching in a dense environment. Trajectories with less than 9 timesteps, corresponding to about 90 − 120 s, are rejected. We calculate velocities using a Savitzky-Golay Filter using a 30 − 45 s window to fit 2nd-order polynomials and take a smooth derivative. The mean-squared displacement (MSD) of individual trajectories is calculated by a time average MSD(*τ*) = ⟨**x**(*t* + *τ*)**x**(*t*)⟩_*t*_ up to a maximal displacement of half the trajectories length (*τ* ≤ *T/*2). Features such as Betti numbers and edge lengths are calculated on the graph. The Euler characteristics are calculated from the mask using region properties of scikit-image [62].

### Supplementary Text I: Ecology and Light fluctuations

Here we briefly discuss the typical light intensities and fluctuations experienced in the natural habitat of *Pyrocystis lunula*. Surface waves typically induce light fluctuations at frequencies of 0.1 − 1 Hz [56], while clouds can cause more pronounced changes in light intensity over the course of minutes [66]. *Pyrocystis spp*. are known to live in the lower eutrophic zone at depths of 60 − 100 m [42] and undergo strong diurnal vertical migration [50]. The growth of *P. lunula* reaches full saturation under low light conditions found at approximately 50 m depth, equivalent to about 12 mW*/*cm^2^ [67]. These natural conditions therefore correspond to the regions in which *P. lunula* adapts its chloroplast extension in response to varying light intensities (Fig. 1D).

### Supplementary Text II: Active Kelvin-Voigt model

To mathematically describe light-induced chloroplast contraction and expansion, we employ a one-dimensional visco-elastic model. This choice is grounded on the uni-axial contraction of the chloroplast network, with negligible expansion or contraction occurring in the perpendicular directions (Fig. 2A,B,D, Movie S4-S6). We assume, that stresses are not externally imposed but applied along the chloroplast network itself. This assumption is supported by the observed structural similarity between actin-network and chloroplast structures [39]. However, assuming constant rheological properties along the network and a linear force transmission, the picture of global stress application or local stresses is equivalent. Viscous dissipation dampens the motion. Here, we assume this dissipation results either from a rheological feature (visco-elasticity) of the contracting network itself, the motion of the chloroplast network through the surrounding viscous cytoplasm, or from a combination of both mechanisms. The precise origin of this feature remains unspecified in our model. Experimental measurements indicate that the fluctuations in chloroplast position are small compared to the global motion (Fig. 1B,D, Movie S1-S3). This allows us to neglect noise in this model. Thus, the stress-induced motion can be formalized as a visco-elastic Kelvin-Voigt solid:

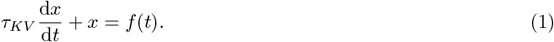

Here *x* is a displacement from an equilibrium position *x*_0_ = 0 and *f* = *F/k* is an equilibrium position of the spring with spring constant *k* forced with *F*(*t*). 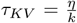 is a visco-elastic relaxation-timescale, where *η* is a viscous damping factor. If light *I*(*t*) is sensed, the contractile force is gradually increased by a chemical concentration *c*: *f* (*t*) = *βc*(*t*) hence the equilibrium position changes over time, until it reaches an intensity-dependent set-point. We model such a first-order reaction by a process with a timescale *τ*_1_ (fig. S2):

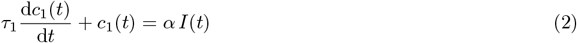

with *α* such that max(*c*) = 1. Moreover, we allow the relaxation timescale to take two different values *τ*_*KV*_ and 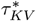, depending on whether light *I* is “on” (*I >* 0) or “off” (*I* = 0)^1^. Therefore, contraction and expansion will follow different timescales, which is likely a direct consequence of different rates of the underlying molecular contractile/extensile mechanism. As we measure *x* from the spread-out position 𝒜*/ 𝒜*_0_ = 1 we define: 𝒜 (*t*)*/ 𝒜*_0_ = 1 − *L* ∗ *x*(*t*) and incorporate *L* in *β* for convenience, without loss of generality. These equations can be solved analytically for constant light irradiation starting at *t* = 0 (as in our experiments Fig. 1):

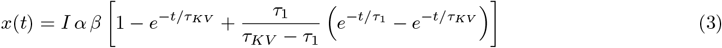

For simplicity we denote Δ𝒜_*max*_ ≡ *αβI*.

#### Transient response at low light *I < I*_*th*_

At low light intensities, we observe an adaptive response (Fig. 1D). Similarly to models of light adaptation in green algae [19], we postulate that below a threshold intensity when 0 *< I < I*_*th*_, a second light sensor sends an opposing signal (fig. S2), which controls the chloroplast spreading (accumulation response). The second sensor is modeled as

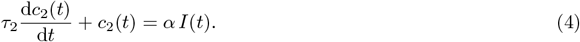

Here the driving force is defined by the difference: *f* = *β*(*c*_1_ − *c*_2_). For times greater than *T*_*max*_ = *argmax*(*x*) the direction of chloroplast motion reverses (expansion) and thus, as the time-scale of this outward-motion was found to be different, equation (5) will then be used with 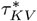. The analytical solution for the case of 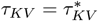 reads:

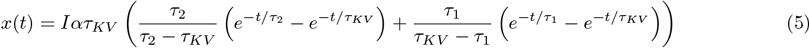

#### Dynamic filter-properties of chloroplast response

The main driving factor of the observed behavior is the optimization of photosynthesis, while minimizing photodamage. We interpret such a response as a result of the cells signal-processing mechanism upon light sensing. Assuming there is a cost to such a response, the relevant light fluctuations have to be filtered out from the irrelevant ones. Consequently, our model can be viewed from the perspective of linear filters (if *I > I*_*th*_). We can easily see that the super-threshold dynamic equations (1) and (4) will restore a harmonic oscillator by inserting equation (1) into (2) with *f* = *τ*_*KV*_ *c*_1_ (simplifying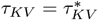):

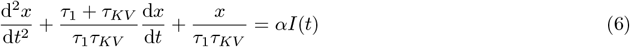

For any combination of the *τ*_*KV*_ and *τ*_1_ this system is over-damped as 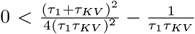^2^. If our light-input is Fourier-transformable, i.e. 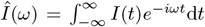 exists, we can find a linear response relationship 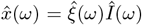, with a susceptibility 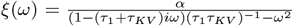, showing that the system essentially behaves as a long-pass filter, cutting off the high frequencies from noisy environmental fluctuations and adapting towards slow trends of light at time scales *τ* ≈ 3 − 5 min.

#### Model fitting

We fit the models to every experiment and report the mean fit parameter and standard deviations. The *R*^2^ score consistently exceeds 0.95, indicating a reasonably good fit, despite the low complexity of the model and the noisy data. The fitting scheme used is a least-squares fit. The fitting parameters remained notably consistent for all different experiments and physiological conditions::

| Figure | Irradiance [mW/cm<sup>2</sup>] | $\tau_1$ [min] | $\tau_2$ [min] | $\tau_{KV}$ [min] | $\tau_{KV}^*$ [min] | $\Delta\mathcal{A}_{max}$ |
| --- | --- | --- | --- | --- | --- | --- |
| Fig. 1B, S4A-D | 2.8 | 1.1 | - | 3.3 | 9 | 0.33 |
| Fig. 1D | 0.6 | $1.6 \pm 0.5$ | $2.5 \pm 0.9$ | $2.9 \pm 0.7$ | $6.2 \pm 2.5$ | $0.40 \pm 0.12$ |
| Fig. 1D | 2.8 | $1.5 \pm 0.7$ | - | $4.4 \pm 0.9$ | - | $0.23 \pm 0.04$ |
| Fig. 1D | 4.9 | $1.8 \pm 0.4$ | - | $3.6 \pm 1.3$ | - | $0.43 \pm 0.03$ |
| Fig. 1D | 7.6 | $1.7 \pm 0.5$ | - | $3.7 \pm 1.1$ | - | $0.43 \pm 0.06$ |
| Fig. 1D | 14.5 | $2.0 \pm 0.3$ | - | $2.2 \pm 1.0$ | - | $0.38 \pm 0.06$ |
| Fig. 1D | 23.4 | $2.1 \pm 0.2$ | - | $2.4 \pm 0.2$ | - | $0.36 \pm 0.05$ |
| Fig. 1D | 32.4 | $1.9 \pm 0.7$ | - | $2.9 \pm 0.6$ | - | $0.43 \pm 0.02$ |
| Fig. 1D | 41.4 | $2.0 \pm 0.2$ | - | $2.3 \pm 0.2$ | - | $0.34 \pm 0.05$ |
| Fig. 3G | 3 | $1.47 \pm 0.02$ | - | $1.47 \pm 0.02$ | - | $0.65 \pm 0.01$ |
| Fig. 4C top | 3.25 | $2.5 \pm 0.5$ | $3.9 \pm 0.5$ | $2.8 \pm 0.5$ | - | 1 |
| Fig. 4C bottom | 3.25 | $2.1 \pm 0.5$ | - | $2.1 \pm 0.5$ | - | $0.53 \pm 0.01$ |
| Fig. 4D top | 3.25 | $1.9 \pm 0.5$ | - | $1.9 \pm 0.5$ | - | $0.80 \pm 0.01$ |
| Fig. 4D bottom | 3.25 | $2.2 \pm 0.5$ | - | $2.2 \pm 0.5$ | - | $0.73 \pm 0.01$ |

<sup>2</sup> $0 < \frac{(\tau_1 + \tau_{KV})^2}{4(\tau_1\tau_{KV})^2} - \frac{1}{\tau_1\tau_{KV}} \Leftrightarrow 0 < \tau_1^2 + 2\tau_1\tau_{KV} + \tau_{KV}^2 - 4\tau_1\tau_{KV} = (\tau_1 - \tau_{KV})^2 \square$

### Supplementary Text III: Pharmacological treatment

Cells were treated with 5 *µ*M Nocodazole (microtubule depolimerization), 10 *µ*M Latrunculin B (actin depolimeization) or 2 mM BDM (myosin inhibition), all prepared in aqueous f/2 solution. Prior to treatment, cells were adapted to either darkness or bright light conditions (*I* ≈ 30 mW*/*cm^2^). Following the treatments, the cells were incubated for another 2 h in darkness, or bright light conditions (*I* ≈ 30 mW*/*cm^2^). Microscopy is performed simultaneously for all treatment groups: first dark- and light-adapted cells are subjected to darkness for 70 min. This was followed by exposure to intermediate light levels, with a subsequent increase in light intensity after 30 min.

To measure the effect, we count the cells and measure the overall increase or decrease of the average pixel values with respect to the first time point. The data is normalized for the number of cells in the field of view, to account for size effects.

The unperturbed control-group of light adapted cells (*N* = 1588) expands at dim light conditions within 50 min (fig. S3A,B). BDM (*N* = 1138) and Nocodazole (*N* = 1239) treated cells do not change and expand as efficiently as the control group. Latrunculin B-treated cells (*N* = 1724) do not expand their chloroplasts, as confirmed by the vanishing slope of the relative normalized absorbance (fig. S3A) and by visual inspection (fig. S3B).

For the group of dark adapted cells, which are subjected to intermediate light intensities (Time ≤ 30 min) and strong light intensities (Time *>* 30 min), we find that all groups but the Latrunculin-B treated cells (*N* = 1724) adapt to the strong light stimulus, as confirmed by the decline in absorbance and visual inspection (fig. S3C,D). Our pharmacological inhibitions clearly suggests that the actin network is used both in chloroplast contraction and expansion. While the role of myosin and microtubules is not fully clear and might be concentration dependent, as suggested in studies on the diurnal chloroplast motion [39].

## SI MOVIES

**Movie S1:** Chloroplast contraction of *P. lunula* at different light intensities: (left) 0.6 mW*/*cm^2^, (center) 2.8 mW*/*cm^2^ and (right) 41.4 mW*/*cm^2^, corresponding to S1A-C, respectively.

**Movie S2:** Dynamic light-controlled chloroplast motion with 10 min 0.4 mW*/*cm^2^ red-light imaging between 2.5 min-lasting white-light stimulation at *I* = 7.6 mW*/*cm^2^.

**Movie S3:** Dynamic light-controlled chloroplast motion with 15 min 0.4 mW*/*cm^2^ red-light imaging between 15 min-lasting white-light stimulation at *I* = 7.6 mW*/*cm^2^.

**Movie S4:** Time series for global stimulation with blue light (470 ± 50 nm. Imaging of chlorophyll auto-fluorescence (red look-up table (LUT)).

**Movie S5:** Time series for a second global stimulation with blue light (470 ± 50 nm. Imaging of chlorophyll auto-fluorescence (red look-up table (LUT)). Sequential buckling of cytoplasmic strands clearly visible.

**Movie S6:** Time series for a global stimulation with blue light (470 ± 50 nm. Imaging of chlorophyll auto-fluorescence (red look-up table (LUT)). After 14 min blue light is switched off and ambient red light is placed. Chloroplasts spread out within a larger time scale. Note the adjusted time step.

**Movie S7:** Network analysis of chloroplast autofluorescence signal. Nodes (dots) between the edges of the skeletonized image are tracked over time. Dataset corresponds to Fig. 3 and Movie S4.

**Movie S8:** Peripheral stimulation 488 nm-laser (white box). Chloroplast auto fluorescence (red LUT).

**Movie S9:** Sequential peripheral stimulation 488 nm-laser (white box). Chloroplast auto fluorescence (red LUT). Chloroplast shrinks first on upper stimulation side, while simultaneously the lower side reacts. Then a second stimulus is applied to the lower side.

**Movie S10:** Central stimulation with 488 nm-laser (white box). Chloroplast auto fluorescence (red LUT).

## SI FIGURES

**Fig. S1:**
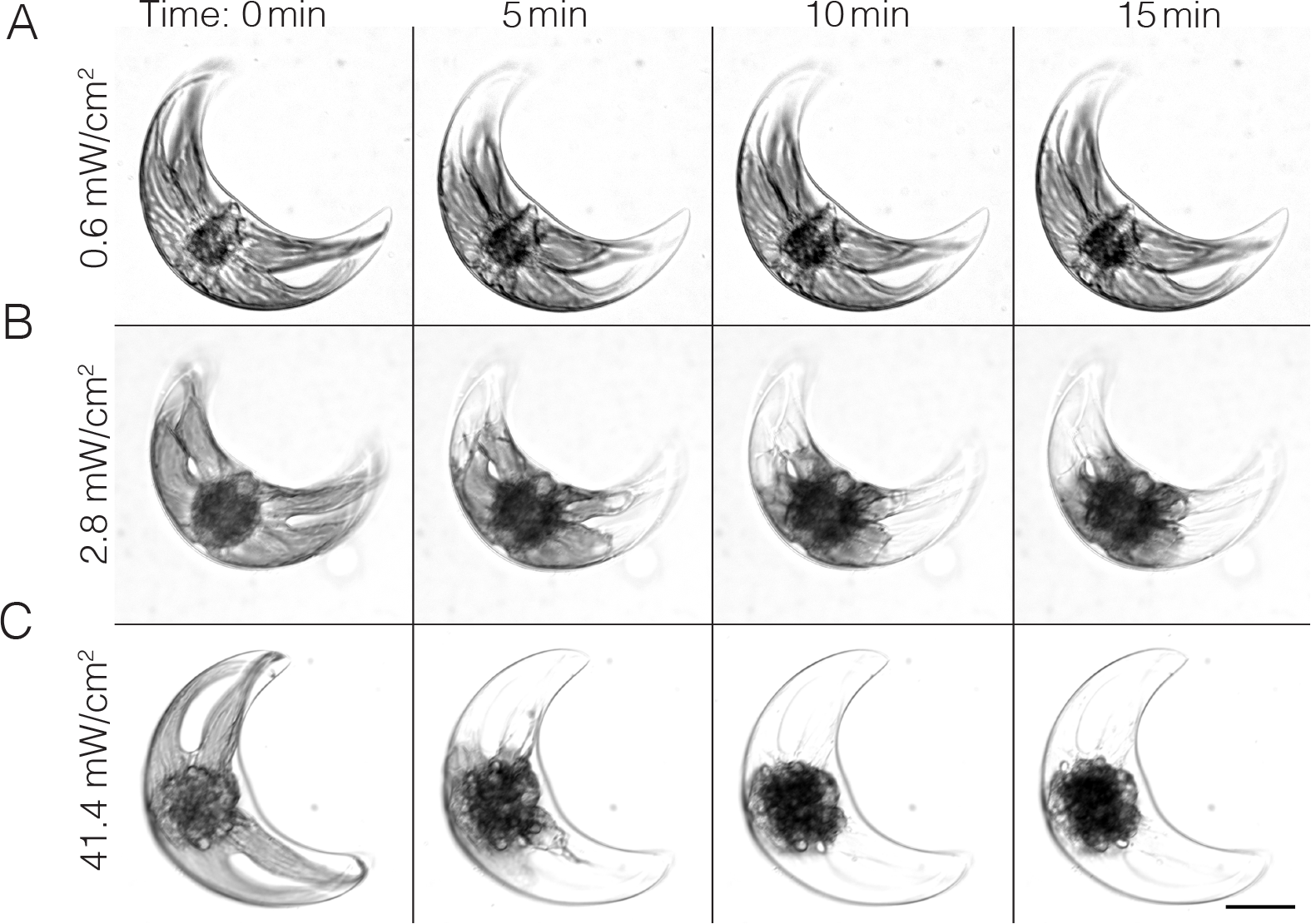
Irradiance-dependent chloroplast retraction. (**A**) at low intensities, chloroplast contraction is limited to a transient response. (**B**) Intermediate intensities lead to an incomplete retraction of the chloroplast. (**C**) High-intensity light stimulation leads to a complete retraction of the chloroplast towards the cytoplasmic core area. Scale bar: 30 *µ*m

**Fig. S2:**
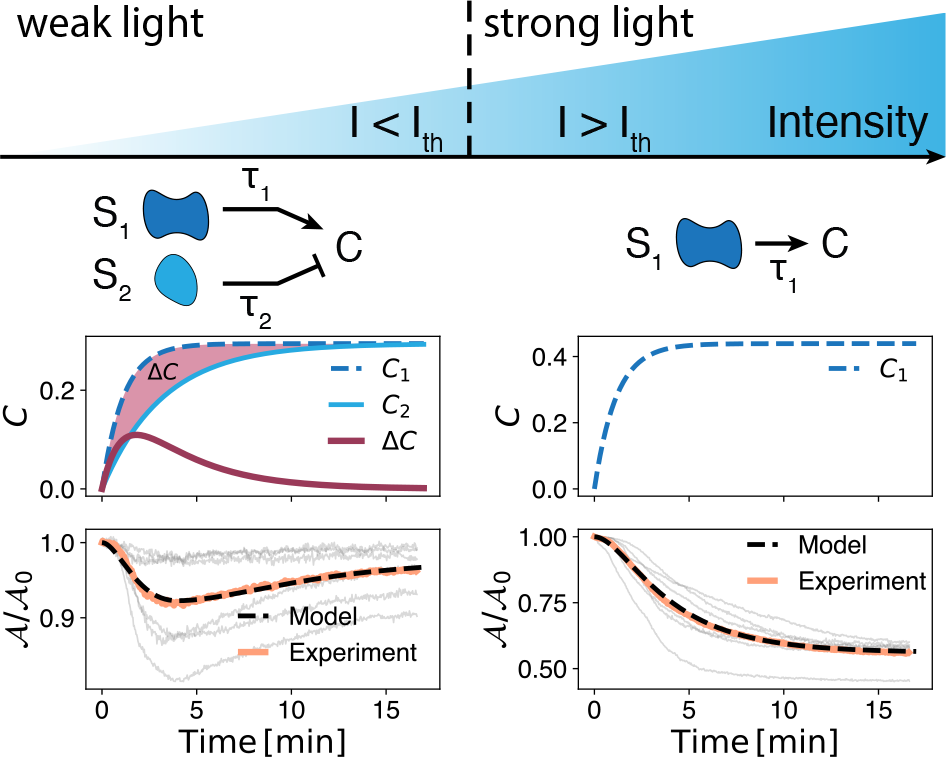
Schematic representation of irradiance-dependent signaling model. Left: for small intensities, light reacts with two opposing light sensors (*S*_1_ and *S*_2_) with two different time scales *τ*_1_ *< τ*_2_, leading to a non-monotonic curve of the concentration Δ*C* (magenta line), and hence to a non-monotonic change in area of the chloroplast (dotted black line, lower graph). As the light intensity is larger than a threshold *I* ≥ *I*_*th*_ the second sensor *S*_2_ is not opposing the signal of sensor *S*_1_ any further. Hence the response is purely photoavoidant (monotonous decrease of chloroplast area 𝒜).

**Fig. S3:**
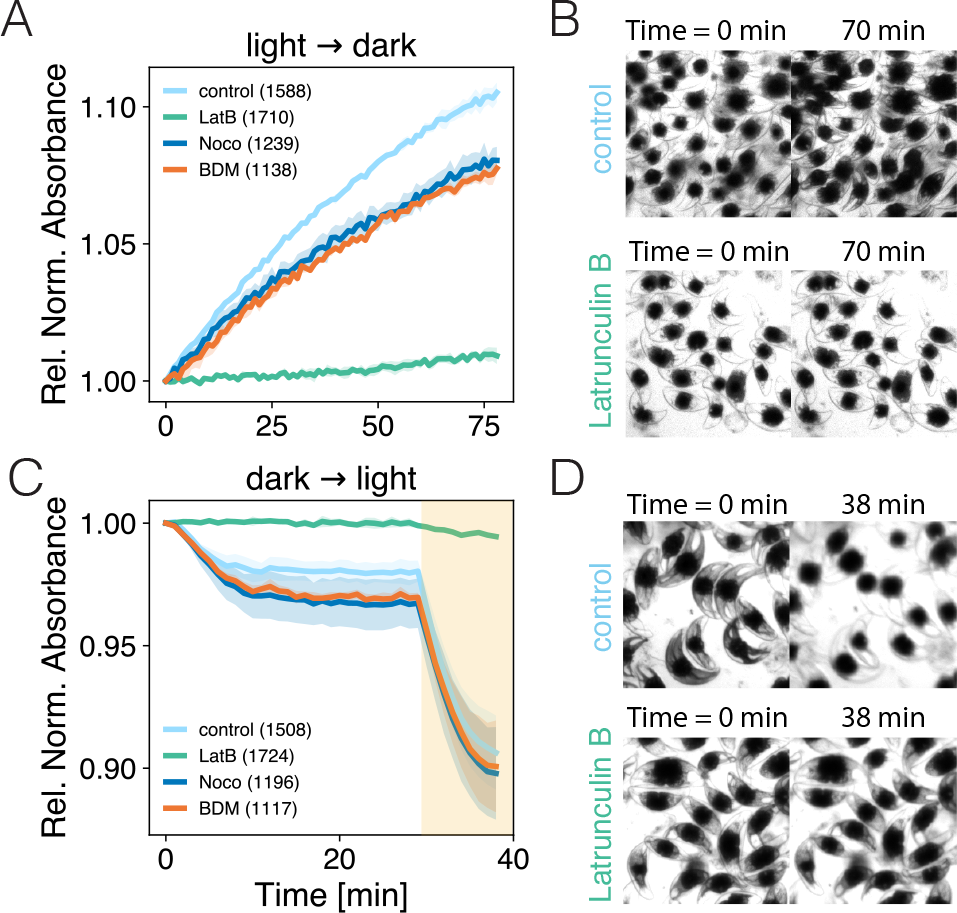
Analysis of pharmacological perturbation to the photoadaptation mechanism. (**A**) Light-adapted cells are placed in dark light conditions and spread out (control). Absorption is measured and normalized with the amount of cells per experiment. 10 *µ*M Latrunculin B treated cells (LatB) do not react. Both 5 *µ*M Nocodazole (Noco) and 2 mM 2,3-butanedione monoxime (BDM) do not show a significant effect. Numbers in the legend stand for the total amount of cells per treatment. Shadowed regions display the min-max error of two independent batches. (**B**) Example images for control and Latrunculin B treated cells before and after dark adaptation. (**C**) Absorption measurement of dim light-adapted cells placed in bright light, with subsequent increase of light intensity (shaded area). (**D**) Example for control and Latrunculin B treated cells before and after bright light adaptation.

**Fig. S4:**
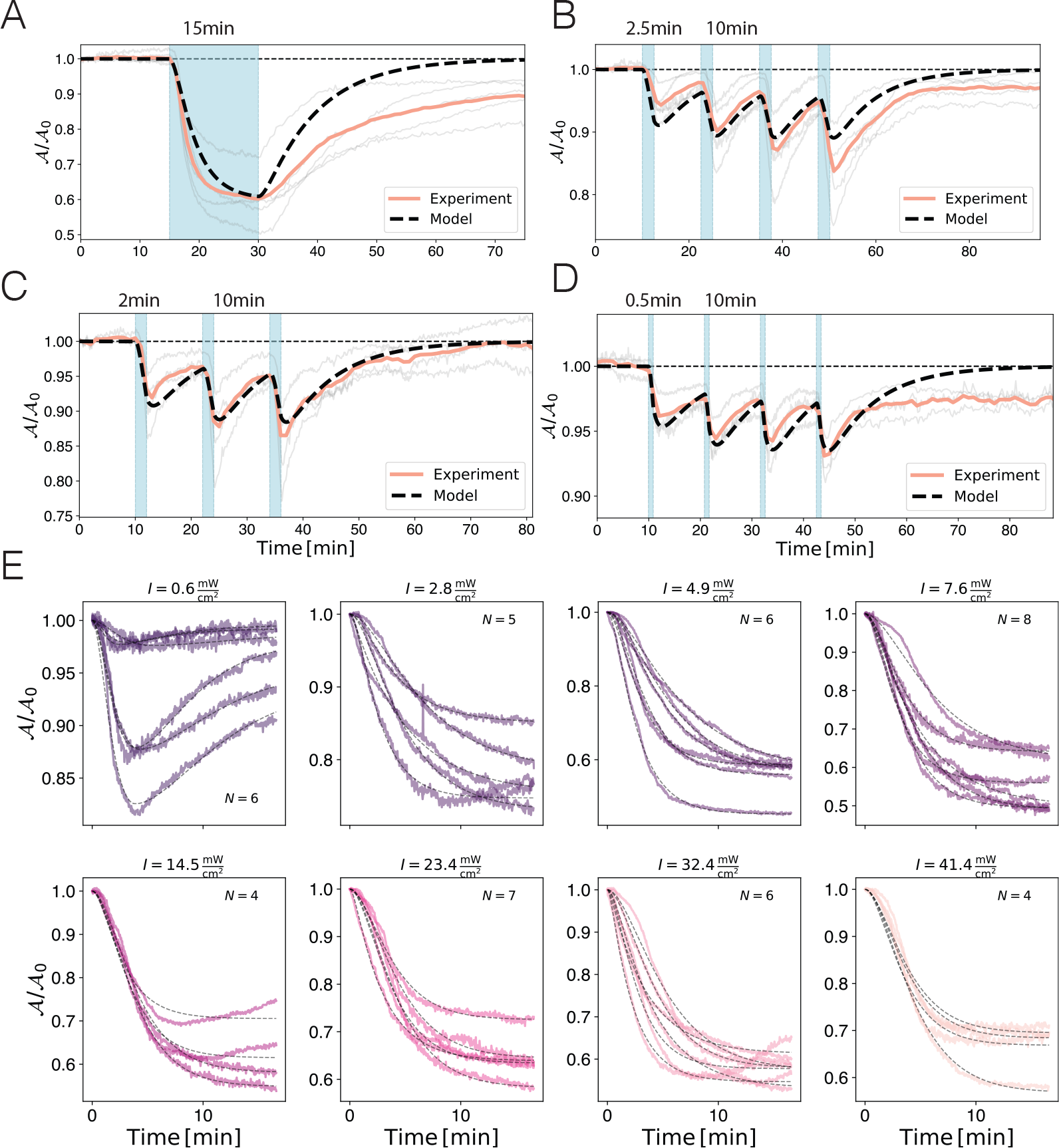
(A-D) Integration of dynamical model (for *I > I*_*th*_) (dotted line) compared to experimental data (orange: mean) for different durations of light illumination (blue region) and dim red light. Parameters for the model were *τ*_1_ = 1.1 min, *τ*_*KV*_ = 3.3 min, 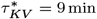 (Table 1). At long times the area was underestimated. (E) Fits of all individual experiments at different light intensities (colors).

## Footnotes

1 Here “on” corresponds to blue or white light stimulation and “off” to dim red light in the experiments.

2 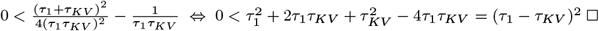

## Notes

### Competing Interest Statement

The authors have declared no competing interest.

